# Optimization for High-Throughput BiFC Screening

**DOI:** 10.1101/2023.10.09.561405

**Authors:** Yunlong Jia, Jonathan Reboulet, Françoise Bleicher, Agnès Dumont, Sylvie Di Ruscio, Benjamin Gillet, Sandrine Hughes, Samir Merabet

## Abstract

The Cell-PCA screen, since its inception, has provided an efficient method for analyzing cellular interactomes and has been used in various biological studies involving proteins like MYC, PER2, and ERK. With rapid advancements in biotechnology, including tools for protein function investigation, the Cell-PCA screen remains relevant. However, despite its successful application in recent studies, there are areas for optimization to ensure its continued relevance in the face of evolving technological advancements.

## Introduction

Since the original Cell-PCA screen was developed and demonstrated a robust and economical high-throughput approach to systematic dissection of cellular interactomes, more recently it has been implemented in a series of projects engaging with many target proteins in various biological contexts, such as MYC, PER2, ERK, etc. Nowadays, biotechnological advances are leaping ahead with new instrumentation and state-of-the-art molecular toolkits. For example, versatile gain-of-function (ORF-overexpression based) and loss-of-function (RNAi or CRISPR/CAS9) screenings have blown the way of functional protein investigation. Though our Cell-PCA screen performed an appropriate readout in different studies *(Delage et al., 2023; Jia et al., 2023)*, even leaky ORF expression existing in the cold CC-ORF cell library, some optimizations are still needed, for not only a known-issue patch, but also aiming to keep Cell-PCA a comparable method alongside of ever-changing technical progress.

## MATERIALS AND METHODS

### Cell line and Plasmids

HEK-293T cells were purchased from Europen Collection of Authenticated Cell Cutures (ECACC) through the biological resource center CelluloNet (AniRA platform of Lyon). Cells were cultured in Dulbecco’s modified Eagle’s medium (DMEM-GlutaMAX-I, Gibco by Life Technologies) supplemented with 10% (v/v) heat inactivated fetal bovine serum (FBS) and 1% (v/v) Penicillin-Streptomycin (5,000U penicillin and 5mg streptomycin/mL), incubating at 37°C, in an atmosphere of 5 % CO2.

The plix-403 lentiviral vector was bought from addgene (plasmid # 41395). This plasmid was subsequently modified by digestion and ligation with MCS (multiple cloning site sequence) oligo. The resulting plix-MCS was further used as backbone of plix-CC-mCherry and plix-CC-PBX1. Plasmid plix-VN-HOXA9 as mentioned in previous part (Part 3.1, Chapter II) was used to generate the plix-VN-HOXA9-PGK-BleoR-mCherry, with PURO element replaced by BleoR sequence. All restriction enzymes were purchased from New England Biolabs. The subsequent ligation reaction were performed by T4 DNA ligase (Promega, France) and transformed into one shot Top10 competent cells (Invitrogen, Cat No. C404010). DNA sequencing were carried out at GENEWIZ Company (Germany). All described vectors are freely available for academic use and can be requested from the authors.

### Transfection

For transfection, 3 × 10^5^ cells were seeded on glass coverslips in 6-well plates. Twenty-four hours after plating, cells were transfected with jetPRIME (Polyplus, Ref 114-15) following manufacturer’s instruction. 1ug of each plasmid DNA were transfected per well, according to different test conditions. After 18 hours of incubation in the presence of doxycycline (100 ng/ml final), the cell-coated coverslip were taken and mounted carefully on a glass slide for image capture under confocal microscopy (Zeiss LSM780). All samples were imaged using identical settings and quantified as previously described *(Dard et al., 2018*, *2019)*.

### Generation of Stable cell line

For stable transfection, HEK-293T cell line was used. The cells were seeded in 6-well plate one day before transfection. 3μg of plix-VN-HOXA9-PGK-BleoR-mCherry construct were transfected, containing the BleoR gene (resistant to Zeocin), using with jetPRIME (Polyplus, Ref 114-15) following manufacturer’s instruction. Seven hours post-transfection, change the transfection medium by new fresh cell growth medium and turn the dish to the incubator. After 40h, check the cell confluence, if cell confluence ≥80%, the selection was initiated with 200μg/ml of Zeocin (Invitrogen, France). Bleo-resistant cells were dominated after 5 days of Zeocin selection and propagated using the same medium, transfer established cells into a big vessel, such as T75 flask. The Zeocin-selection lasted for about 3weeks. Final polyclonal stable cell line was cryopreserved in liquid nitrogen for further usage.

### Functional titration by qPCR

A Biorad CFX96 (Biorad, France) was used for all qPCR measurements. To each reaction (10 μl) containing 5ul iTaq™ Universal SYBR® Green Supermix (2x) (BioRad) and 500 nM of each primer,

WPRE [reaction 1] :

*Forward: ACAATTCCGTGGTGTTGTCG*

*Reverse: AAGGGACGTAGCAGAAGGAC*

RPPH1 [reaction 2]:

*Forward: AATGGGCGGAGGAGAGTAGT*

*Reverse: AGCTTGGAACAGACTCACGG*

Plus, we added 4 μl diluted genomic or plasmid DNA (50ng). The temperature profile was 95°C for 3 min followed by 40 cycles of amplification (95°C for 5s, 60°C for 30 s). All samples were analyzed by melting curve analysis (65-95°C, 0.5°C increments at 2-5 s/step). Vector copy numbers in cells are normalized to human RPPH1 gene copies and presented as proviral copies per genome equivalent. Calculate titers (Transdution units per ml, TU ml^-1^) according to the following formula:

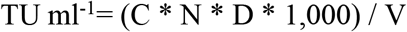

where C = proviral copies per genome, N = number of cells at time of transduction, D = dilution of vector preparation, V = volume (ul) of diluted vector added in each well for transduction. The detailed protocol is available on request.

### New protocol of pooled plasmid DNA preparation and transformation

The protocol of pooled plasmid DNA preparation was provided by the Genetic Perturbation Platform (GPP) (https://portals.broadinstitute.org/gpp/public/resources/protocols).

The collection of 1837 TF ORFs was divided in 7 minipools of average 262 hORFs (the true ORF number varies from 140 to 362). For each minipool, the hORFs were cloned *en masse* from the pDONR223 into the lentiviral expression vector plix-CC-Gateway by Gateway LR reaction (Invitrogen, CAT. 11791020) following manufacturer’s instruction.

To select for the successfully cloned CC-ORFs, each reaction mixtures were transformed into Endura™ electro-competent cells (Lucigen, CAT. 60242-1) and transformed bacteria were selected on LB + Ampicillin plates. Finally, collect all bacterial colonies by scraping plate and proceed to lentiviral plix-CC-TF plasmid extraction using one maxi-prep reaction for each minipool.

## RESULTS

### Design and Performance of new lentiviral vector

Firstly, given the leaky TRE (Tet Response Element) promoter of lentiviral vector pLV-CC-ORF, the 2^nd^ generation Tet-On (Tet-Advanced) system was used in the new lentiviral vector, plix-403. To assess the performance of this new vector, the reporter mCherry was used to constitute plix-CC-mCherry and pLV-CC-mCherry, respectively (**Figure 1A**). The basal expression of mCherry was evaluated by transfection of HEK-293 cells in a Dox-free complete media. In Tet-On system, herein plix-CC-mCherry, new transactivator rtTA (reverse tetracycline-controlled transactivator) was created by fusing rTetR with VP16, which reversed the phenotype and created a reliance on the presence of tetracycline for induction, rather than repression. The rtTA will not bind TRE promoter to induce the gene expression, in absence of Dox. In contrast, pLV-CC-mCherry belongs to part of the orignal first generation of Tet-Off Systems, which is tTA (tetracycline-controlled transactivator)-dependent to promote expression. The result demonstrated that pLV-CC-mCherry has proximately 7-fold higher basal expression than that of plix-CC-mCherry (**Figure 1B**). New lentiviral plasmid plix-403 offered a significant improvement over the original first generation doxycycline-inducible system, such as previous pLV, with significantly reduced basal expression, mitigating the adverse effect from potential genetic perturbations.

**Figure 1.**
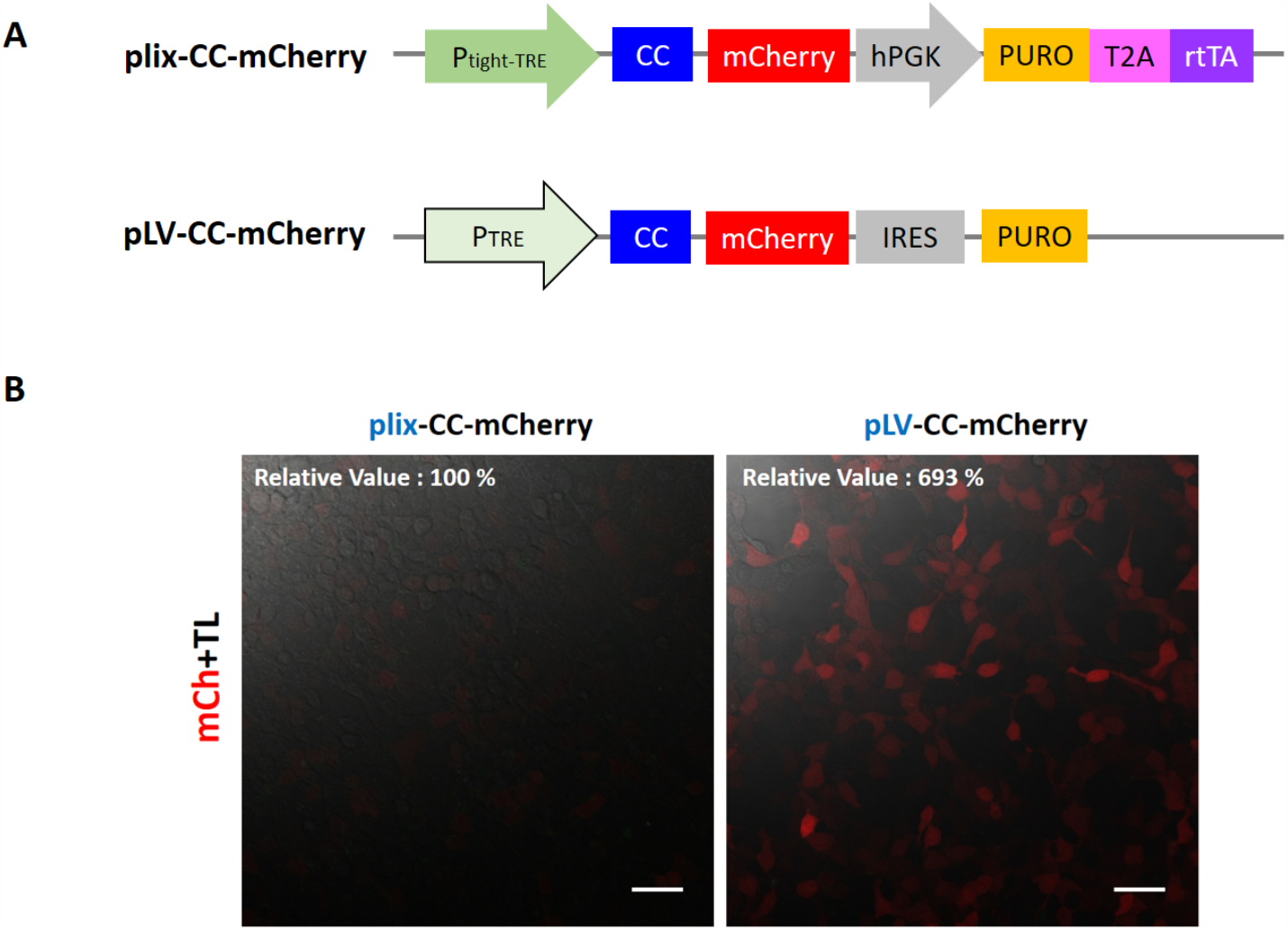
(A) Illustration of plix-CC-mCherry and pLV-CC-mCherry constructs. P_tight-TRE_, Tet-responsive tight promoter as 2^nd^ generation doxycycline-inducible version, consisting of seven tet operator sequences followed by the minimal CMV promoter. P_TRE_, original Tet-responsive promoter. CC, C-terminal fragment of mCerulean, 155-238aa. hPGK, human phosphoglycerate kinase 1 promoter. IRES, internal Ribosome Entry Site, allows for initiation of translation from an internal region of the mRNA. PURO, puromycin-resistant element. T2A, self-cleaving 2A peptides. **(B) Comparison of basal fluorescent signals between plix-CC-mCherry and pLV-CC-mCherry in HEK cells**. Images were taken by Zeiss confocal microscope at x20 objective. mCh, channel mCherry. TL, channel transmitted light. Scale bar = 50 μm. (Jonathan Reboulet, unpublished data)

Moreover, in new plix-403 plasmid, hPGK promotor confers an independent moderate expression of PURO, which is favorable to select and maintain the established stable cell line without Dox, eliminating the further mutual influence with puromycine during a long-term culture. Another advantage of plix-403 is using a 3rd-generation vectors for lentivirus packaging system, in which including a chimeric 5’LTR removes the requirement for the HIV Tat protein, thus HIV Tat protein, thus decreasing the probability of creating replication-competent lentivirus in your target cells with a lower biosafety risk.

Accordingly, we decided to replace the pLV-with plix-403-based cell library preparation. Therefore, to know the plix-403 performance at different concentration of Dox is important when using this Tet-On system. Concentration gradient experiment was successful to prove the sensitivity and rigorousness of plix-based system (**Figure 2**). Henceforward, the transfected gene expression can be easily adjusted by Dox concentration as different test conditions need.

**Figure 2.**
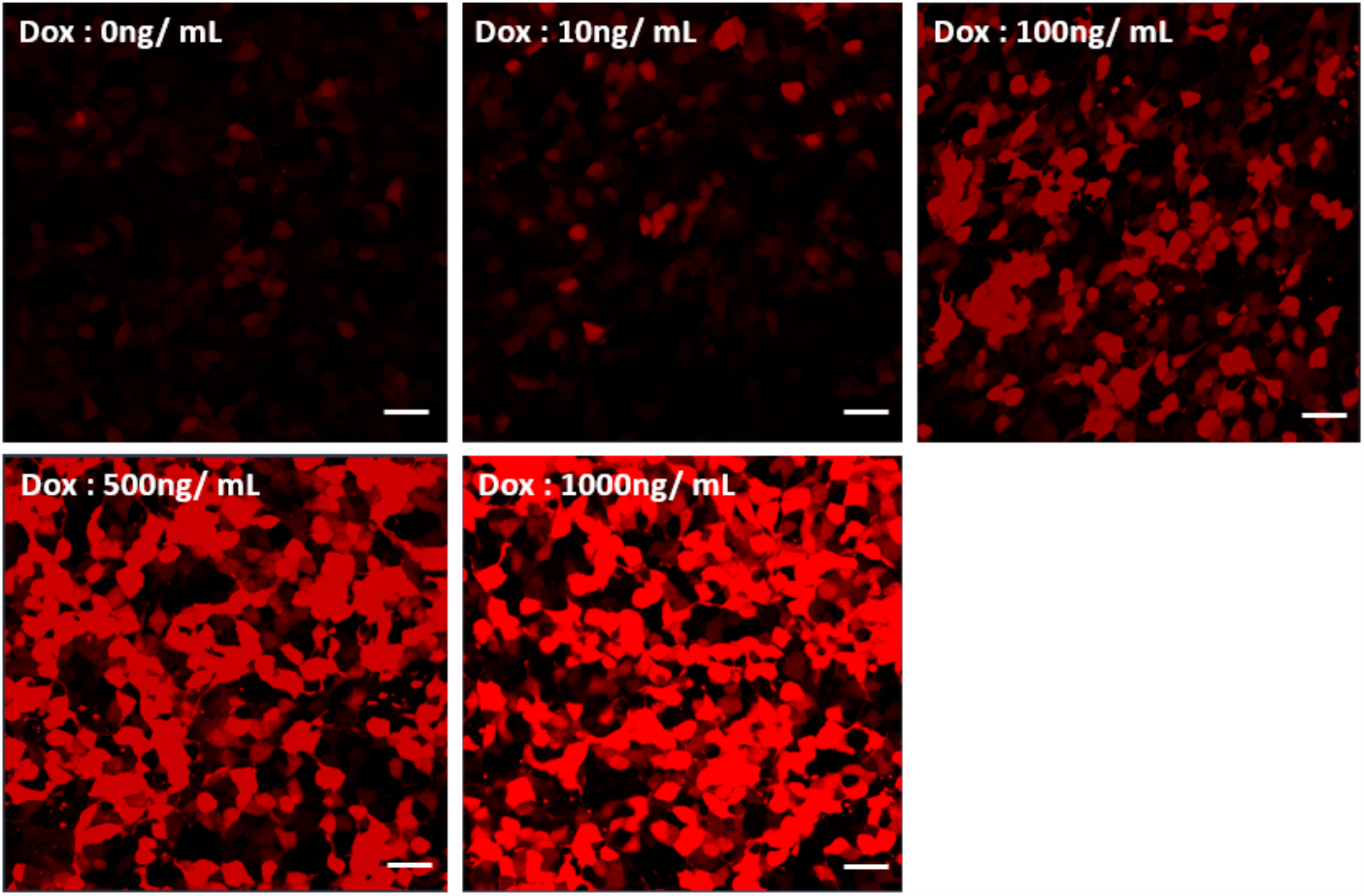
Testing Tet-On system using plix-CC-mCherry. HEK cells were transfected by plix-CC-mCherry and incubated with complete media containing 0 to 1000 ng/mL Dox for 18 h. Images were taken by Zeiss confocal microscope at x20 objective. Scale bar = 50 μm.

The plix-based plasmid, concerning BiFC, can be used for both VN-bait and CC-prey proteins. For optimizing the final BiFC-positive cell sorting and collection, reporter fluorescent protein mCherry was chosen as transfection marker. The proof-of-principle test was carried out on VN-fused HOXA9 bait protein (**Figure 3A**). The plix-VN-HOXA9-mCherry plasmid was first tested by transfection with plix-CC-PBX1, which showed a good colocalization profile for BiFC fluorescence signals and mCherry, indicating all the VN-HOXA9/CC-PBX1 reassembly co-occurred with expression of mCherry marker (**Figure 3B**). Furthermore, the attempting stable cell line harboring endogenous mCherry marker was established by plix-VN-HOXA9-mCherry. The single CC-PBX1 prey was tested by transfection and the BiFC assay displayed clear and pertinent protein interaction loci as with double transfection system (**Figure 3C**), implying the feasibility of future transfection-free BiFC screening (BleoR is secondary antibiotic selection marker, working independently of PURO).

**Figure 3.**
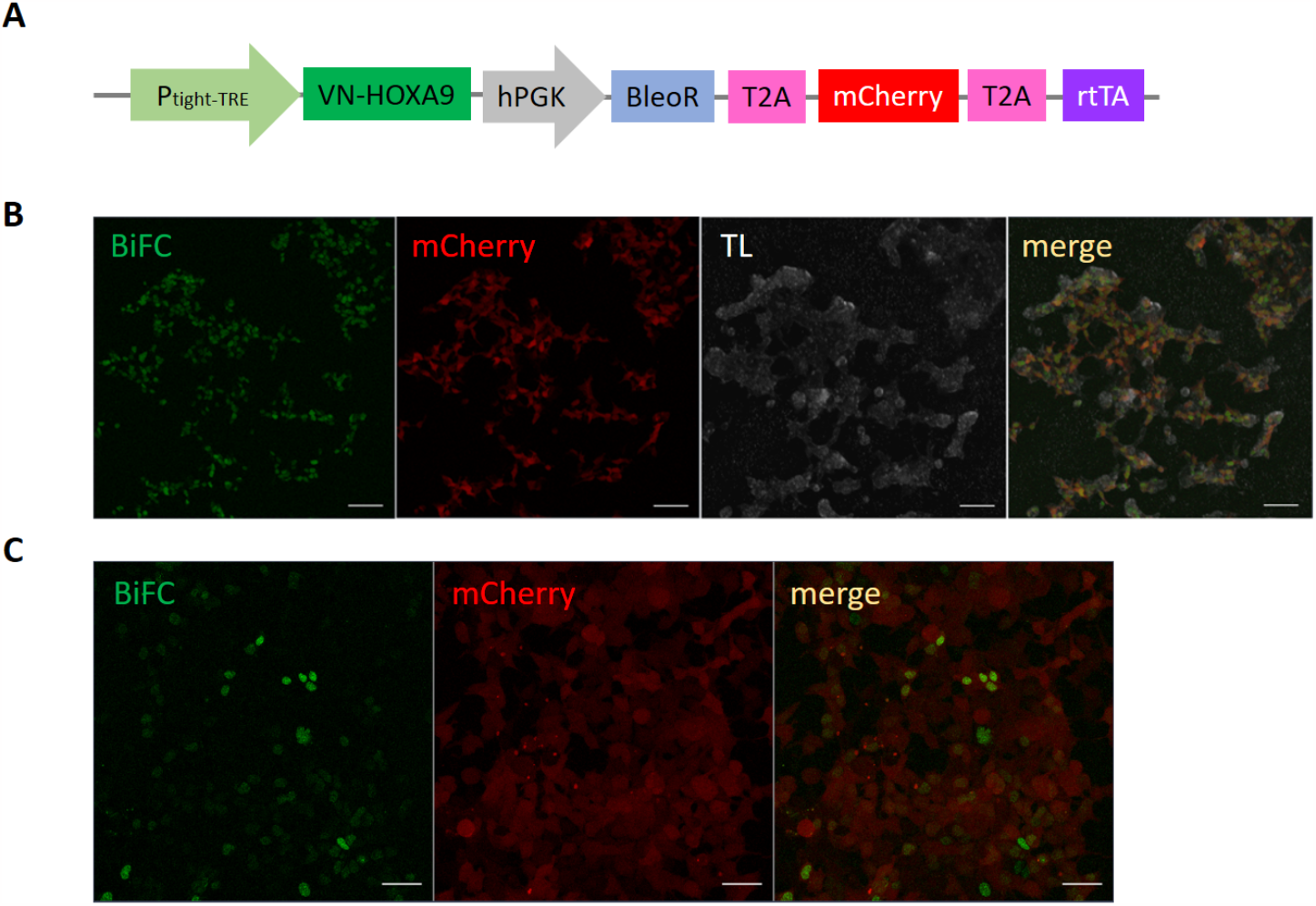
Construction and performance of plix-VN-HOXA9-mCherry. (A) Illustration of plix-VN-HOXA9-hPGK-BleoR-mCherry. BleoR, confers resistance to bleomycin, phleomycin, and Zeocin, (B) BiFC performance test of plix-VN-HOXA9-mCherry cotransfected with plix-CC-PBX1. The test was executed in live HEK cells at Dox concentration of 100ng/mL. Images were taken by Zeiss confocal microscope at x10 objective. Scale bar = 100 μm. (C) BiFC test in plix-VN-HOXA9-mCherry stable cell line. One plasmid transfection was performed by plix-CC-PBX1 in live cells at Dox concentration of 100ng/ml. Images were taken by Zeiss confocal microscope at x20 objective. Scale bar = 50 μm.

### Lentivirus functional titration

Functional titers measure how many viral particles can infect your target cells, which is critical for subsequent one-ORF-copy-to-one-cell infection. The popular FACS-based titration is only available when viral vectors carrying fluorescent markers. Consequently, our previous pLV-CC-ORF lentiviral library titer was measured by antibiotic-dependent colony counting method. However, it may underestimate viral titer and can not handle the multiple integration events. In addition, the experiments are time-consuming and prone to large inaccuracy. As such, the most accurate method of titration, qPCR, was applied to the plix-CC-ORF lentiviral library. For example of functional titration by qPCR, the result from one newly made plix-CC-ORF pooled lentivirus was shown in **Figure 4**. The absolute copy numbers of WPRE and RPPH1 were calculated based on standard curve (**Figure 4A**), in a range of virus dilutions (from 0ul to 80ul). The final functional titer can vary from different diluted conditions (TU, transduction units; from 5.48 X 10^6^ to 1.23 X 10^7^ TU/mL) (**Figure 4B**, right). The acceptable titer should meet these criteria: the calculated titer should be similar between different DNA quantities in the same dilution condition; the ratio of maximal and minimal titer should not exceed five. Either max or min titer could be used for subsequent stable cell line establishing procedure, but the MOI (Multiplicity of Infection) and cell number may be adjusted according to your final titer choosing. Moreover, the virus copy number per cell is another important information, which could be obtained by this qPCR titration method (**Figure 4B**, left).

**Figure 4.**
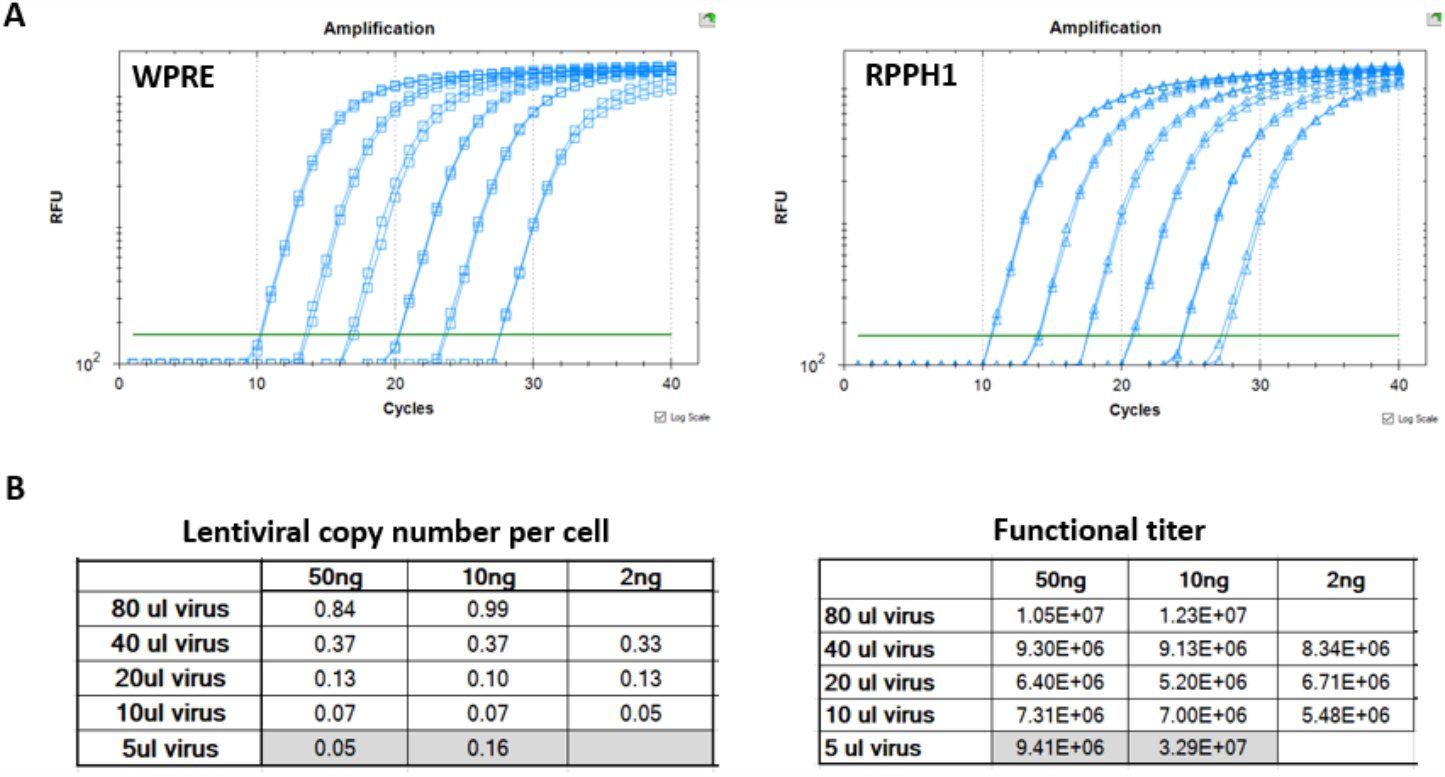
Functional titration of plix-CC-ORF viral library by qPCR. (A) Standard curve of WPRE and RPPH1. WPRE, Woodchuck Hepatitis Virus (WHV) Posttranscriptional Regulatory Element, as marker gene of lentivirus for copy number determination. RPPH1, the RNA component of the RNase P ribonucleoprotein, a gene that exists as a single copy per haploid genome (or 2 copies per human cell). (B) Quantification of lentiviral copy number per cell and functional titer. The values in grey cells are discarded, due to big discrepancies.

#### 3.2.3.3 Custom sub-library: hORFeome v8.1-based human transcription factor library

Complete sets of cloned protein-encoding open reading frames (ORFs), as known as ORFeomes, are essential tools for large-scale proteomics and systems biology studies. Human ORFeome clone collection is currently the largest publicly available resource of full-length human ORFs, which is created by the OC (http://www.orfeomecollaboration.org/). To achieve a nearly entire set of ORFs, expansion of hORFeome library is still ongoing, for example the latest version v9.1 containing 17,408 protein-coding genes *(Luck et al., 2020)*. Likewise, a recent update was made for our previous hORFeome stock, which was now replaced by hORFeome v8.1 purchased from BioCat GmbH. This collection represents almost 12,000 unique genes, and NGS-confirmed ORFs as a set of Gateway Entry clones ready for transfer to Gateway-compatible expression vectors, assigning high accuracy and quality to this bacterial glycerol stock. Based on this new hORFeome, a sublibrary comprising all known human TFs was made by a multi-functional robotic platform. There are two reasons to undertake this sublibrary. First, TFs are of the most interest in biological community as potential research subjects of various gene function-related studies. The screening dedicated to known TFs is highly needed. Second, the new protocols of cloning and pooled plasmid DNA production were used, which will further improve the ORF representation in final pooled library and enable a less biased deep sequencing.

The known TFs were manually curated, referring to two previous studies *(Chawla et al., 2013; Lambert et al., 2018)*. 1837 unique TF ORFs were cherry-picked from whole hORFeome v8.1 glycerol stock by high-throughput robotic liquid handling (**Figure 5A**). For the sake of more efficient pooled transformation, new electro-competent cells were used, which fulfilled transformant colony number at least 1000x greater than the number of constructs in the pooled library (**Figure 5B**).

**Figure 5.**
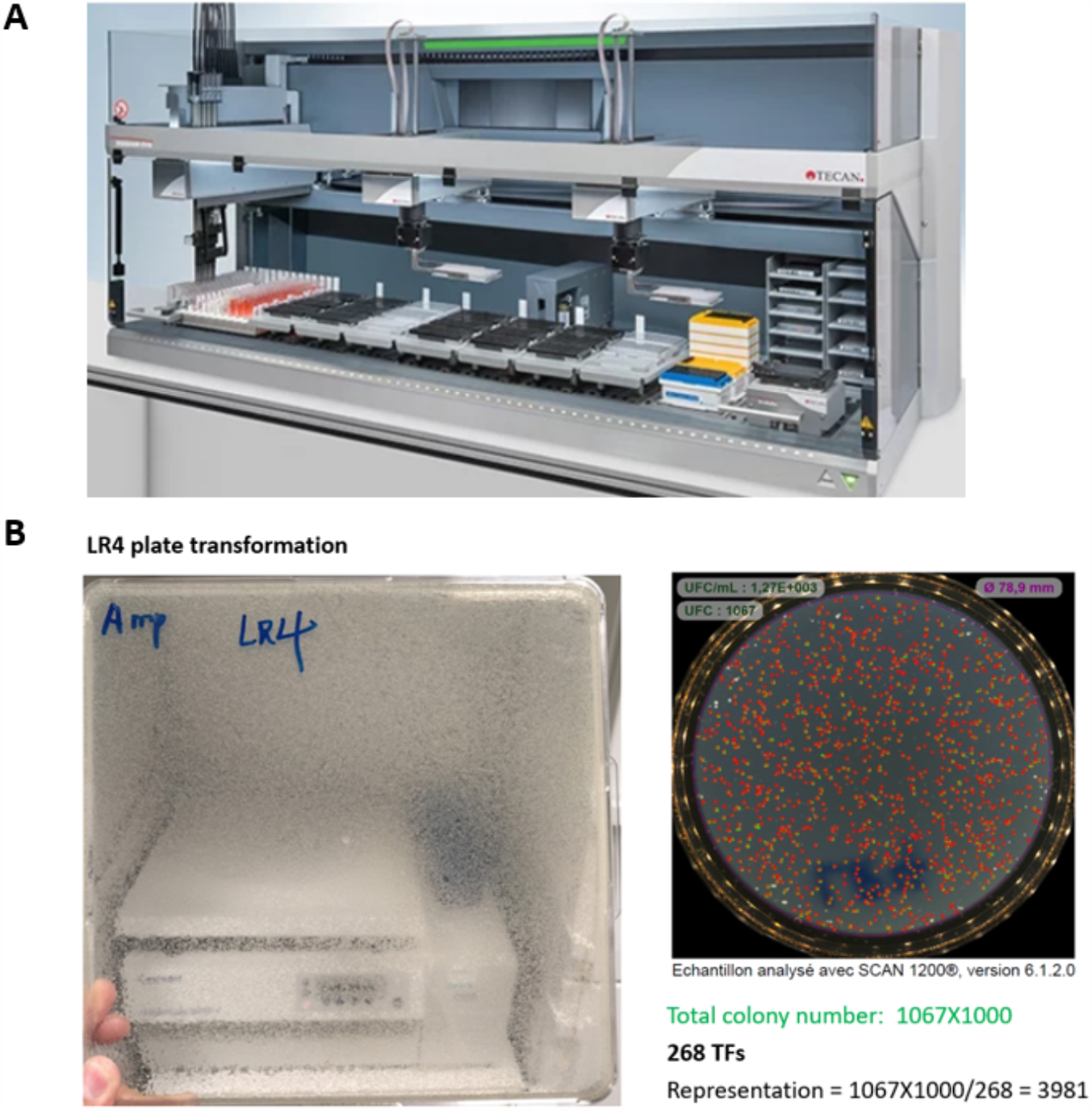
(A) Multi-functional robotic platform, FREEDOM EVO 200. The largest workstation in the Freedom EVO series, it provides an extensive work area and variable configurations, with a choice of liquid handling and robotic arms. **(B) New transformation performance, illustrated by 268 TFs-minipool transformation**. 1ul of deactivated Gateway LR reaction used in new competent cells according to the manufacturer’s instructions. The transformant was plated on a square Bioassay dish (left). The resulting colony number was calculated by petri dish colony counting, on which contains 1/1000 of original transformant. The representation was calculated attaining outperformed X3881.

## DISCUSSION

Apart from some of the representative results that were given above, there are still other optimizations addressing our BiFC screening strategy. For instance, the illumina NextSeq 500 system was used in our ORF deconvolution instead of ever Ion Torrent PGM sequencing, which enables a highly-multiplexing barcode sequencing with a much greater depth. Benefited from this new NGS platform, our new plix-CC-ORF pooled plasmid was sequenced in a full plasmid manner, as it was previously reported that this strategy yielded smaller coverage variability (**Figure 6**) *(Yang et al., 2011)*. Furthermore, to alleviate the side effect of cell death during FACS that largely affects the sorted population purity and limits the cell sorting speed, one reagent, named CellCover, is undergoing testing. It claimed that this reagent is compatible with all cell fixation-based downstream applications, without chemical crosslinking, including FACS and (single cell-based) NGS, protein sequencing, and more. However, it is worth noting that not one single method assesses all the specific and true positive PPIs, thus application of orthogonal methods is highly important to chart a more accurate PPI readout. For this purpose, co-IP, as the gold standard assay for PPIs, is considered to be performed as a complementary method for HT-BiFC putative candidates. Owing to recent technical advances, Jess automates traditional Western blotting while maximizing multiplexing with multiple detection channels, also compatible for a mid-throughput co-IP assay. Truly, automation of protein separation and immune-detection eliminates many of the tedious, error-prone steps of traditional blotting that limit data quality. This system is recently available in our region (Protein Science Facility, SFR Biosciences, Lyon), which is worth trying in the next step.

**Figure 6.**
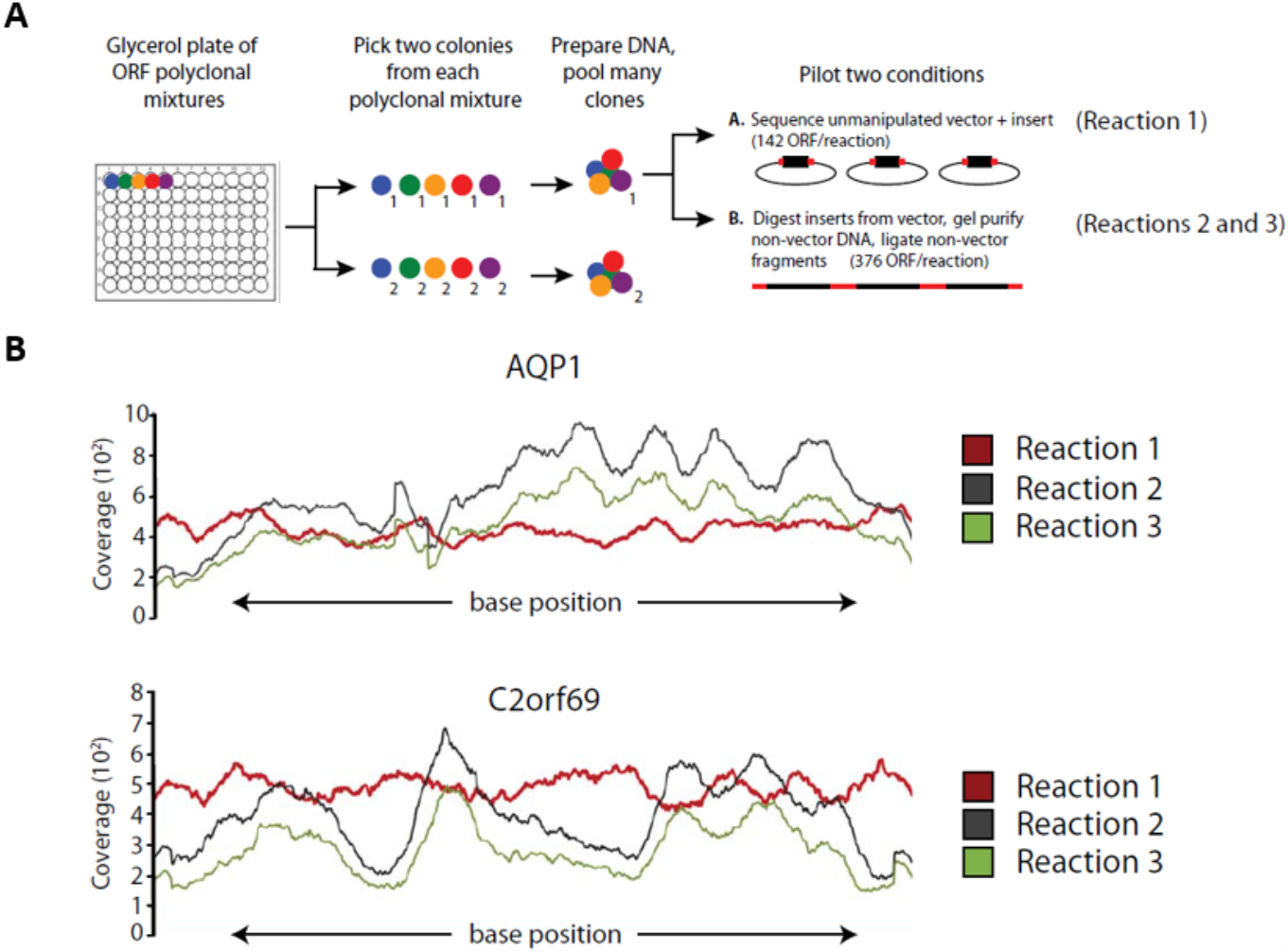
Pilot experiments to optimize pooling strategy for next generation sequencing of ORF clones. (A) Schematic of pilot. Two conditions were tested. Reaction 1 consisted of 142 unmanipulated ORF plasmids. Reactions 2-3 were duplicate reactions in which ORF inserts from 376 ORF plasmids were enzymatically purified from plasmid backbones. (B) Coverage across the length of 2 representative ORFs. Adapted from Supplementary figure 1 in *(Yang et al., 2011)*.

Reflecting on the optimized screening and to-do list, collectively, a highly-improved HT-BiFC screening method is expected in the near future, which will undoubtedly contribute to the endeavor of promising rigorous and robotic multi-omics study.

## Notes

### Competing Interest Statement

The authors have declared no competing interest.

